# Improved high-quality genome assembly and annotation of Tibetan hulless barley

**DOI:** 10.1101/409136

**Authors:** Xingquan Zeng, Xiangfeng Li, Lijun Bai, Yulin Wang, Tong Xu, Hongjun Yuan, Qijun Xu, Sang Zha, Zexiu Wei, Wangmu Qimei, Yuzhen Basang, Jiabu Dunzhu, Mingzai Yu, Tashi Nyima

## Abstract

**Background:** The Tibetan hulless barley (Hordeum vulgare L. var. nudum), also called “Qingke” in Chinese and “Ne” in Tibetan, is the staple food for Tibetans and an important livestock feed in the Tibetan Plateau. The Tibetan hulless barley in China has about 3500 years of cultivation history, mainly produced in Tibet, Qinghai, Sichuan, Yunnan and other areas. In addition, Tibetan hulless barley has rich nutritional value and outstanding health effects, including the beta glucan, dietary fiber, amylopectin, the contents of trace elements, which are higher than any other cereal crops.

**Findings:** Here, we reported an improved high-quality assembly of Tibetan hulless barley genome with 4.0 Gb in size. We employed the falcon assembly package, scaffolding and error correction tools to finish improvement using PacBio long reads sequencing technology, with contig and scaffold N50 lengths of 1.563Mb and 4.006Mb, respectively, representing more continuous than the original Tibetan hulless barley genome nearly two orders of magnitude. We also re-annotated the new assembly, and reported 61,303 stringent confident putative protein-coding genes, of which 40,457 is HC genes. We have developed a new Tibetan hulless barley genome database (THBGD) to download and use friendly, as well as to better manage the information of the Tibetan hulless barley genetic resources.

**Conclusions:** The availability of new Tibetan hulless barley genome and annotations will take the genetics of Tibetan hulless barley to a new level and will greatly simplify the breeders effort. It will also enrich the granary of the Tibetan people.

## Introduction

Tibetan hulless barley (Hordeum vulgare L. var. nudum) is a cereal crop, a variant of barley, alias called hulless barley, yuan wheat, rice barley(**Fig.1**). Tibetan hulless barley has a long history of cultivation in the Qinghai-Tibet Plateau. The adaptation to extreme environmental conditions in high altitudes made it the staple food for Tibetans beginning at least 3,500–4,000 years ago [1]. It continues to be the predominant crop in Tibet, occupying ∼70% of crop lands.

**Figure 1.**
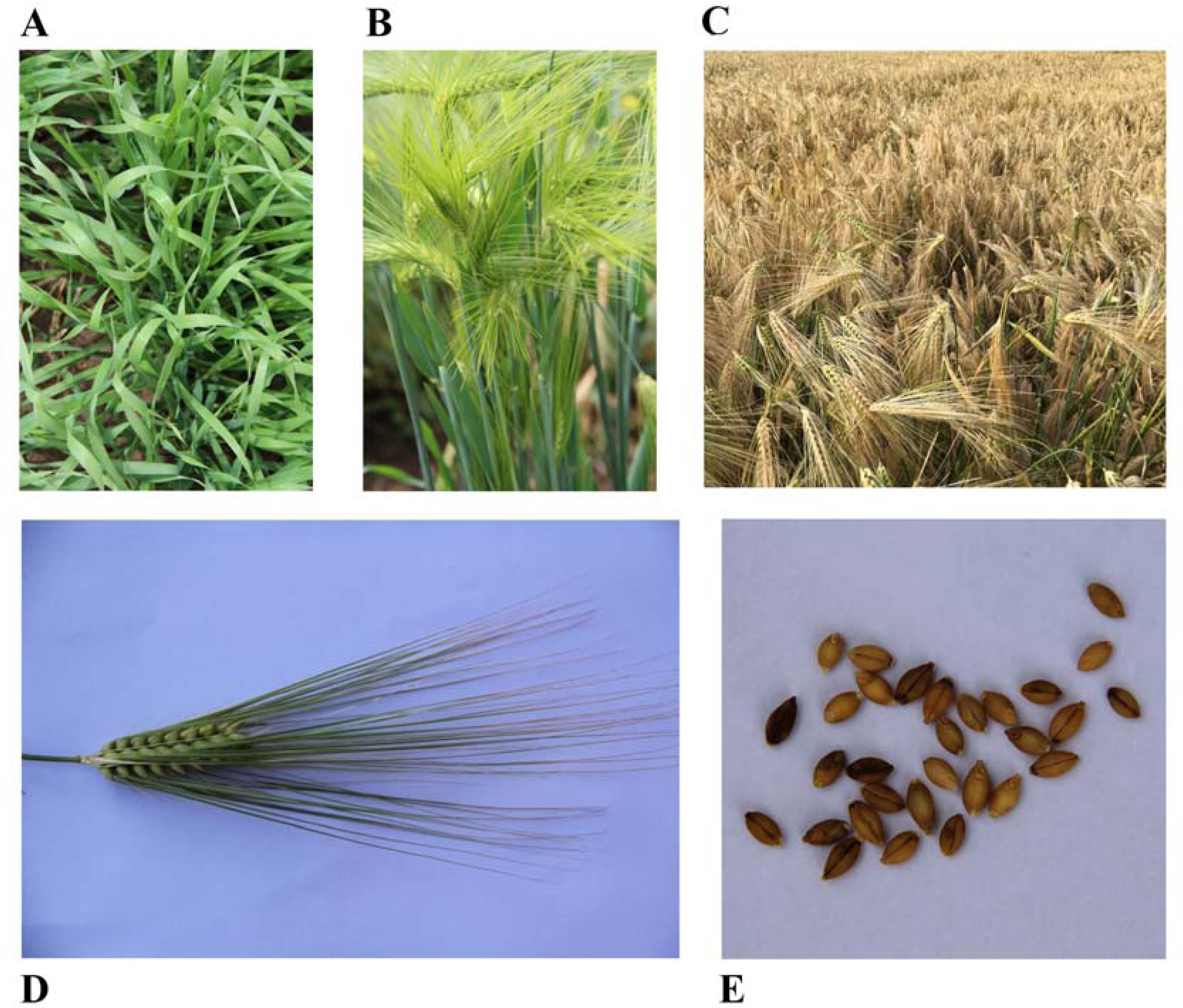
Morphological characteristics of the Tibetan hulless barley, Tibetan hulless barley. These pictures show (A),seedling;(B), heading stage; (C), mature stage;(D), filling stage spike and (E), grain of Tibetan hulless barley.

Genome sequences provide a substantial basis for understanding the biological essences of crops associated with biologically and economically essential traits[2]. Therefore, it is necessary and crucial to construct the genome of a species. In recent years, with the development of NGS sequencing technology, many crops have published the genome draft or near-finished genome sequences, especially the larger genome of the wheat family[3-13]. At the same time, two studies have published two versions of the Tibetan hulless barley genome in 2015[2] and 2017[14], respectively. And the former assembly strain is Lasa Goumang, the latter is Zangqing320, all excellent cultivars of Tibetan hulless barley. The estimated genome size of the Tibetan hulless barley is 4.48Gb[2], which is much smaller than the 5.1Gb[4, 10] of barley. Although the former is a subspecies of barley, considering that the barley genome is rich in repeats(about 80.8%- 84%[2, 4, 10]), the difference in genome size may come from the numbers and length of transposons.

Although the publication of the Tibetan hulless barley genome has attracted a new upsurge in Tibetan hulless barley genetics research[15, 16], due to the current poor genome assembly, such as the particularly short contig, the very low contig N50, and the poor genome annotation, the functional genome studying of Tibetan hulless barley has caused some vital problems. In addition, with the development of single-molecule sequencing platforms, such as those by Pacific Biosciences and Oxford Nanopore Technologies, have become available to researchers and are currently being increasingly applied in *de novo* genome assembly and genome update, and other scaffolding techniques(such as BioNano optical mapping, Chicago, Hi-C and 10X genomics)[17-19], it provides a new opportunity for improving Tibetan hulless barley genome assembly. Therefore, we *de novo* assembled a high-quality genome of Tibetan hulless barley using high-coverage Pacbio long reads, combining published Illumina paired-ends and mate-pairs reads. And then, we applied PacBio Iso-seq to get high confidence full-length transcripts and re-annotation Tibetan hulless barley protein-coding genes. This new assembly, annotations and newly online Tibetan hulless barley genomics database discussed below will serve as valuable resources for investigating the economic and genetic of Tibetan hulless barley and will also help the molecular breeders to continually improve Tibetan hulless barley or barley germplasm resources.

## DNA and RNA sequencing

### DNA isolation, libraries construction and sequencing

The seedling of Tibetan hulless barley (NCBI BioSample ID **SAMN09914874**) were collected from Tibet, China. After collection, tissues were immediately transferred into liquid nitrogen and stored until DNA extraction. DNA was extracted using the Cetyltrimethyl Ammonium Bromide (CTAB) method according to the protocal “ Preparing Arabidopsis Genomic DNA for Size-Selected ∼20 kb SMRTbell^™^ Libraries ”(www.pacb.com/wp-content/uploads/2015/09/Shared-Protocol-Preparing-Arabidopsis-DNA-for-20-kb-SMRTbell-Libraries.pdf). The quality of the extracted genomic DNA was checked using 1% agarose gel electrophoresis, and the concentration was quantified using a Qubit fluorimeter (Invitrogen, Carlsbad, CA, USA).

Single-molecule Real-time long reads sequencing was performed at NextOmics Technology Corporation (Wuhan, China) with a PacBio Sequel Sequencer (Pacific Biosciences, Menlo Park, CA,USA). The SMRT Bell library was prepared using a DNA Template Prep Kit 1.0, and Six 20-kb SMRT Bell libraries were constructed(**Fig. S1**). Genomic DNA(10ug)was mechanically sheared using a Covaris g-Tube with a goal of DNA fragments of approximately 20kb. A Bioanalyzer 2100 12K DNA Chip assay was used to assess the fragment size distribution. Sheared genomics DNA(5ug) was DNA-damage repaired and end-repaired using polishing enzymes. A blunt-end ligation reaction followed by exonuclease treatment was conducted to generate the SMRT Bell template. A Blue Pippin device(Sage Science) was used to size select the SMRT Bell template and enrich large fragments(>10kb).The size-selected library was quality inspected and quantified on an Agilent Bioanalyzer 12Kb DNA Chip (Agilent) and a Qubit fluorimeter (Life Technologies). A ready-to-sequence SMRT Bell Polymerase Complex was created using a Binding Kit 2.0 (PacBio p/n 100-862-200), according to the manufacturer’s instructions. The PacBio Sequel instrument was programmed to load and sequenced the sample on PacBio SMRT cells v2.0 (Pacific Biosciences), acquiring one movie of 360 min per SMRT cell. The MagBead loading (Pacific Biosciences) method was used to improve the enrichment of the larger fragments. A total of 64 SMRT cells were processed yielding 300.24G subread sequences, average 4.69Gb per cell, and 9,358bp in length(**Table S1**).

For Illumina sequencing data, we applied previously published paired-ends and mate-pair library data[2](SRA no. SRA201388, totally 819.37Gb clean data) for PacBio assembly error evaluation and scaffolding, respectively.

### RNA isolation and sequencing

Fresh roots, and leaves of Tibetan hulless barley were sampled in Tibet, China. Totally, 12 samples, different N condition (Prepared for unpublished paper) were used for Iso-seq sequencing. Total RNA was prepared by grinding tissue in TRIzol reagent (Invitrogen 15596026) on dry ice and processed following the protocol provided by the manufacturer. To remove DNA, an aliquot of total RNA was treated with RQ1 DNase (Promega M6101), followed by phenol/chloroform/isoamyl alcohol extraction, chloroform/isoamyl alcohol extraction using Phase Lock Gel Light tubes (5 PRIME 2302800) and ethanol precipitation. Precipitated RNA was stored at -20°C.

One microgram of total RNA per reaction per tissue was reverse transcribed using the Clontech SMARTer cDNA synthesis kit. Specific barcoded oligo dT in separate PCR tubes, to generate barcoded FL cDNA. Large-scale PCR products were purified with AMPure PB beads and quality control (QC) was performed on a 2100 BioAnalyzer (Agilent). Equimolar ratios of six cDNA libraries were pooled together. A total of 3.8 mg cDNA was subjected to size fractionation using the SageELF system. Size fractions eluted from the run were subjected QC and pooled in equimolar ratios for subsequent re-amplification to yield four libraries (1–2, 2–3, 3–6 and 5–10Kb). The pooled PCR products were purified using AMPure PB beads. One to five micrograms of purified amplicons were subjected to Iso-Seq SMRT Bell library preparation (https://pacbio.secure.force.com/SamplePrep). A total of 17 SMRT cells (unpublished data) were sequenced on the PacBio RS II platform using P6-C4 chemistry with 3–4 h movies.

## Genome assembly

The Sequel raw bam files were converted into subreads in fasta format using the standard Pacbio software package BAM2fastx. Totally, 300.24Gb PacBio data, covered genome ∼67X(estimated genome size 4.48Gb),was used to initially assemble.

### Falcon assembly

we used falcon software package (https://github.com/PacificBiosciences/falcon) to construct primary assembly. Firstly, corrected the error reads by the overlap strategy (parameter length_cutoff control is used to correct the wrong seed reads length);Secondly, pre-assembled the corrected reads; Thirdly, Constructed contig with long reads obtained from pre-assembly (parameter length_cutoff_pr controls the pre-assembled reads length used to build contig); After many attempts, we determined that the following two reference settings can get better assembly results: length_cutoff = 5000;length_cutoff_pr = 10000. And, we get the initial assembly, 1.229Mb in length of contig N50.

## Arrow polish

There are often errors in the preliminary assembly results of the genome, such as SNP or InDel, which can be fixed by certain algorithms. Arrow is a genomic polishing tool based on hidden Markov models. We use the following steps to polish the genome:

1) Align the PacBio sequencing reads to the genome assembly with blasr (pbalign --algorithm=blasr, the rest of the parameters default);

2) Use Arrow to polish based on the alignment result (variantCaller -- algorithm=arrow, the rest of the parameters default).pbalign and variantCaller are from the SMRT Link 4 toolkit (https://www.pacb.com/support/software-downloads/)

## Scaffolding

The Scaffolding tool can group, sort, and orient contigs to join contigs into longer sequence fragments, called scaffold. The sequence between some contigs may be unknown but the distance between them can be roughly known, which is also helpful for subsequent related work.

SSPACE[20] is a widely used tool for building scaffold. Firstly aligned the reads to contigs using alignment tools (such as BWA, bowtie), and then built scaffold based on the connection information between the contigs. This project uses Illumina data previously published and PacBio data to build scaffold.

Firstly, using Illumina mate-pair library (20kb and 40Kb fragment long) sequencing data to scaffolding contigs. Aligned the paired-end reads to contigs with bowtie[21]; then, constructed scaffold (parameter: -p 1 -g 2) with SSPACE-STANDARD-3.0.

Secondly, using PacBio data to continuously improving the scaffolds. Aligned PacBio reads to the scaffold version of Illumina data using blasr; and then, built scaffold with SSPACE-LongRead[22].

The final assembly produced 1,856 scaffolds, 1,092 of which greater than or equal to 1Mb, with a contig N50 length of 1.563Mb and scaffold N50 of 4.006Mb. Compared to the preliminary draft of the published two Tibetan hulless barley genomes[2, 14], the contiguity was improved more than ∼80 and 243 times, respectively(**Table 1**;**Table S2- S3;Fig. S2**).

**Table 1.** Comparison of the new genome with previously published assemblies of the Tibetan hulless barley genome.

|  | **This study** | **Zeng *et al.*[2]** | **Dai *et al.*[14]** |
| --- | --- | --- | --- |
| **Sequenced genome size (Gb)** | 4.00 | 3.89 | 3.73 |
| **Contig N50 (Kb)** | 1,563.00 | 18.07 | 5.94 |
| **Scaffold N50 (Kb)** | 4,006.00 | 242.00 | 171.11 |
| **Repeat proportion(%)** | 87.48 | 81.39 | NA |
| **Annotated protein-coding genes** | 61,303* | 36,151 | 46,787 |

\* Only refer to stringent confident putative protein-coding genes.

### Assembly evaluation

First, the 682.57Gb Illumina paired-ends reads(library fragment 250,500 and 800,respectively) were mapped to new assembly, and 99.87-99.89% PE reads were mapped concordantly (**Table S4**).

Using these short reads, the estimated quality value (QV) of Tibetan hulless barley genome was calculated according to a previously described method[23]. The erroneous bases in the genome assembly were identified using the variant calling software Genome Analysis Toolkit (GATK) v4[24] with default parameters. The QV of the Tibetan hulless barley genome was estimated to be 52(**Table S5**), which means that the accuracy of the assembly in base level is very accurate after base correction.

We randomly downloaded 4 *Hordeum vulgare subsp. vulgare* (domesticated barley) BAC clone sequencing data (NCBI ID: AC248476.1,AC250041.1,AC253057.1 and AC253422.1) to evaluate assembly quality. Using nucmer[25] mapping BAC data to new assembly, the results manifested very high consistency.(**Fig. S3**).

In addition, we also used BUSCO[26] to perform an assessment of genome completeness. BUSCO used a total of 1440 genes to evaluate the genome; the genome has 1312 complete BUSCO genes, accounting for 91.1%, of which 1288 have single copies, accounting for 89.4%, 24 have multiple copies, accounting for 1.7%; 34 gene have only a few fragments of the gene; 94 genes are missing from the genome.

Together, the results indicated that our dataset represented a genome assembly with a high level of coverage, quality and completeness.

## Repetitive sequences annotation

Repetitive DNA accounts for much of the remarkable variation in plant genome sizes[27], especially some Gramineae crop plants, above 80% [10, 12, 28] of genome for repetitive elements. Repetitive sequences are divided into transposons and simple repeats. According to Tandem repeats finder program[29],we marked simple repeat elements, 155,813,415 in length,3.89% of genome. For transposons, we performed homology-based and de novo prediction, respectively. The software packages involved are RepeatMasker[30], RepeatProteinMask[30], repeatmodeler (http://www.repeatmasker.org/RepeatModeler/) and ltr_finder[31]. The repetitive database used here is Repbase 21.01[32](**Fig. S4**). Long terminal repeat(LTR) retroelements, belong to class I elements, is accounted for 70.22% of genome. All repeats together accounted for 87.48% of the genome(**Table 2**;**Table S6**).

**Table 2.**
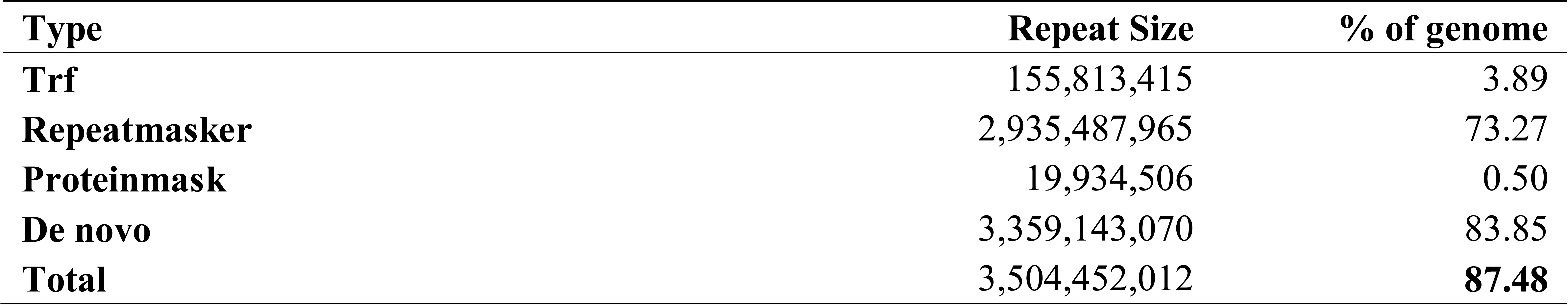
Repetitive element annotation statistics.

## Gene prediction

For the annotation of protein-coding gene, we used a combination of multiple data sources, followed by a high confidence (HC) and low confidence (LC) classification of the annotated genes(**Fig. S4-5**).

### Combination of multiple annotations

According to Iso-seq methods, we obtained 39,442 high-quality full-length transcripts. Theses high-quality transcripts were aligned to the genome using GMAP (version 2017-05-08)[33], with a total of 14,099 perfect alignments; TransDecoder v4.0.1(https://github.com/TransDecoder) was used to predict the coding region of 14,099 transcripts, of which 13,936 contained open reading frame (ORF), these ORF contained transcripts are ultimately assigned to 9,360 genes, called perfect genes. We employed Augustus v3.3[34],which used perfect genes as input train sets, to get 128,400 putative genes. At the same time, we downloaded representative proteins of 7 related species (*Triticum aestivum A*[12]*, Triticum aestivum D*[9], *Brachypodium distachyon*[35], *Hordeum vulgare*[10], *Oryza sativa*[36], *Sorghum bicolor*[37] and *Zea mays*[38]) from public database(**Table S7**). Aligning the protein sequences of each species to the Tibetan hulless barley genome using tblastn v2.6.0+[39]; and then, using Genewise v2.4.1[40] to predict gene structure and obtain homologous gene sequences from various species in the Tibetan hulless barley genome. In addition, using 16 transcriptome data (**Table S8**) from different tissues and different conditions, we mapped the reads to Tibetan hulless barley genome by hisat2 v2.1.0[41] and reference-based assembled the 47,490 transcripts by stringtie v1.2.4[42, 43]. The gene information extracted from PacBio Iso-seq, Augustus, Homolog and RNA-seq was integrated using EVidenceModeler[44], according to different weight parameters (**Table S9**), to obtain 129,269 putative genes. The same RNA-seq reads were de novo assembled using Trinity v2.4.0[45] to obtain 722,803 transcripts. Structural optimization of EVM integration results was performed using PASA v2.1.0[46] and Trinity assembly results.

129,269 transcripts were annotated with NR, Uniprot, and Pfam database. And filter the putative genes according to the function assignment result: (1) NR, Uniprot comparison identity>=50%, coverage>=50%; (2) Not satisfying (1), but protein length >= 50, and containing the Pfam domain. The tandem duplicated genes were identified using MCScanX. Only the longest protein of each subgroup was kept to represent the corresponding gene. The combine Tibetan hulless barley genome annotation included 65,833 genes with names derived from BLAST analysis and genes with domain names derived from Pfam analysis.

### High-confidence Genes

Stringent confidence classification was applied to all predicted genes to discriminate between loci representing high-confidence (HC) protein-coding genes and less reliable low-confidence (LC) genes, which potentially consisted of gene fragments, putative pseudogenes and non-(protein)-coding transcripts. We assigned confidence values to a gene model in a two-step procedure by using the similar criteria and methods described previously[10](**Fig. S5**). Finally, we obtained 61,303 stringent confident putative genes, of which 40,457 is HC genes(**Table 3**; **Fig. S5**).

**Table 3.** Protein-coding genes prediction summary.

| **Method** | **Software and gene set** | **Gene number** |
| --- | --- | --- |
| PacBio | Smrtlink | 39,442 |
|  | Gmap | 14,099 |
|  | TransDecoder | 13,936 |
| Ab initio | Augustus | 128,400 |
| Homology-based | Triticum aestivum A | 42,601 |
|  | Brachypodium distachyon | 33,911 |
|  | Hordeum vulgare | 46,312 |
|  | Oryza sativa | 38,426 |
|  | Sorghum bicolor | 29,909 |
|  | Triticum aestivum D | 44,897 |
|  | Zea mays | 29,872 |
| RNA-seq | StringTie | 47,490 |
| Integration | EvidenceModeler | 129,269 |
|  | PASA | 129,269 |
| Filter | NR、SwissProt、PFAM | 68,018 |
|  | After MCScanX moved | 65,833 |
|  | After TE moved | **61,303** |

Functional annotation of all 61,303 protein sequences was performed using NR, KEGG, SwissProt, Trembl, GO, PFAM, and InterPro databases. 96.57% of all protein sequences were assigned to at least one database(**Table S10**).

### Non-coding Gene prediction

In the process of annotation of non-coding RNAs, tRNAscan-SE[47] software is used to search for tRNA sequences in the genome according to the structural characteristics of tRNA; since rRNA is highly conserved, the rRNA sequence of related species can be selected as a reference sequence, through BLASTN Alignment to find rRNA in the genome; in addition, using the Rfam family’s covariance model, Rfam’s own INFERNAL software[48] can predict miRNA and snRNA sequence information on the genome(**Table 4**).

**Table 4.**
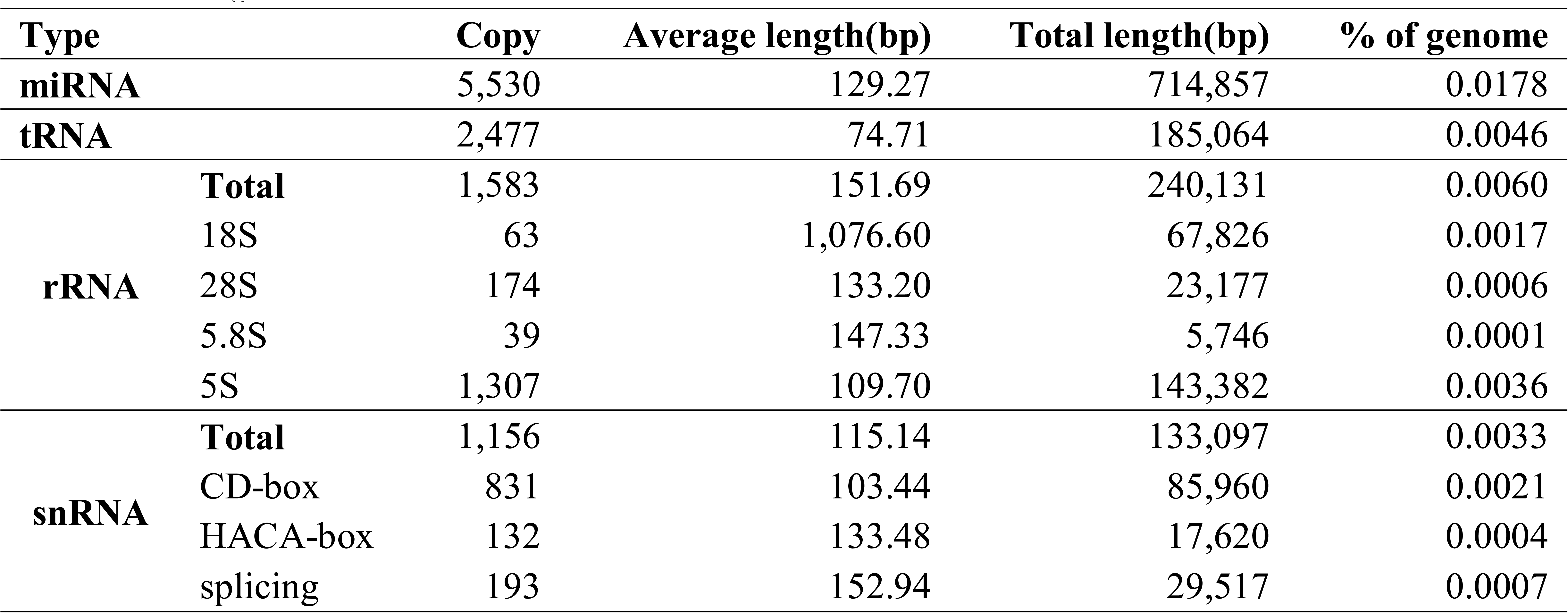
ncRNA genes annotation statistics.

### Annotation evaluation

The BUSCO[26] assessment showed that 91% of the 1,440 genes were completed annotated. And RNA-seq reads rates mapping to gene set also have a significant improvement, from 60% previously annotation to exceed 70%(**Table S8**).

In summary, we also acquired a high-quality genome annotation, and it will greatly benefit future function genomics research.

## Genome database

A new Tibetan hulless barley Genome Database(THBGD) has been developed to facilitate studies on Qinghai-Tibet Plateau crop plant genomics. The database currently provides access to Tibetan hulless barley genome assembly V1,de novo assembly using NGS sequencing and V2,de novo assembly improvement using combined TGS and NGS. Meanwhile, it also provides access to genome annotation, including repetitive element annotation, ncRNA prediction and protein-coding genes prediction(**Fig. 2**). The THBGD embedded JBrowse[49] visual genomic display module to display genome and gene annotation information. One can click on the gene name on the screen to display specific information about the gene. One also can use online tools Blast v2.4.0+ to enter the query sequence in fasta format, align and return the visual aligned results. In the future, we will focus on optimizing the interactive operation and visualization of the THBGD.

**Figure 2.**
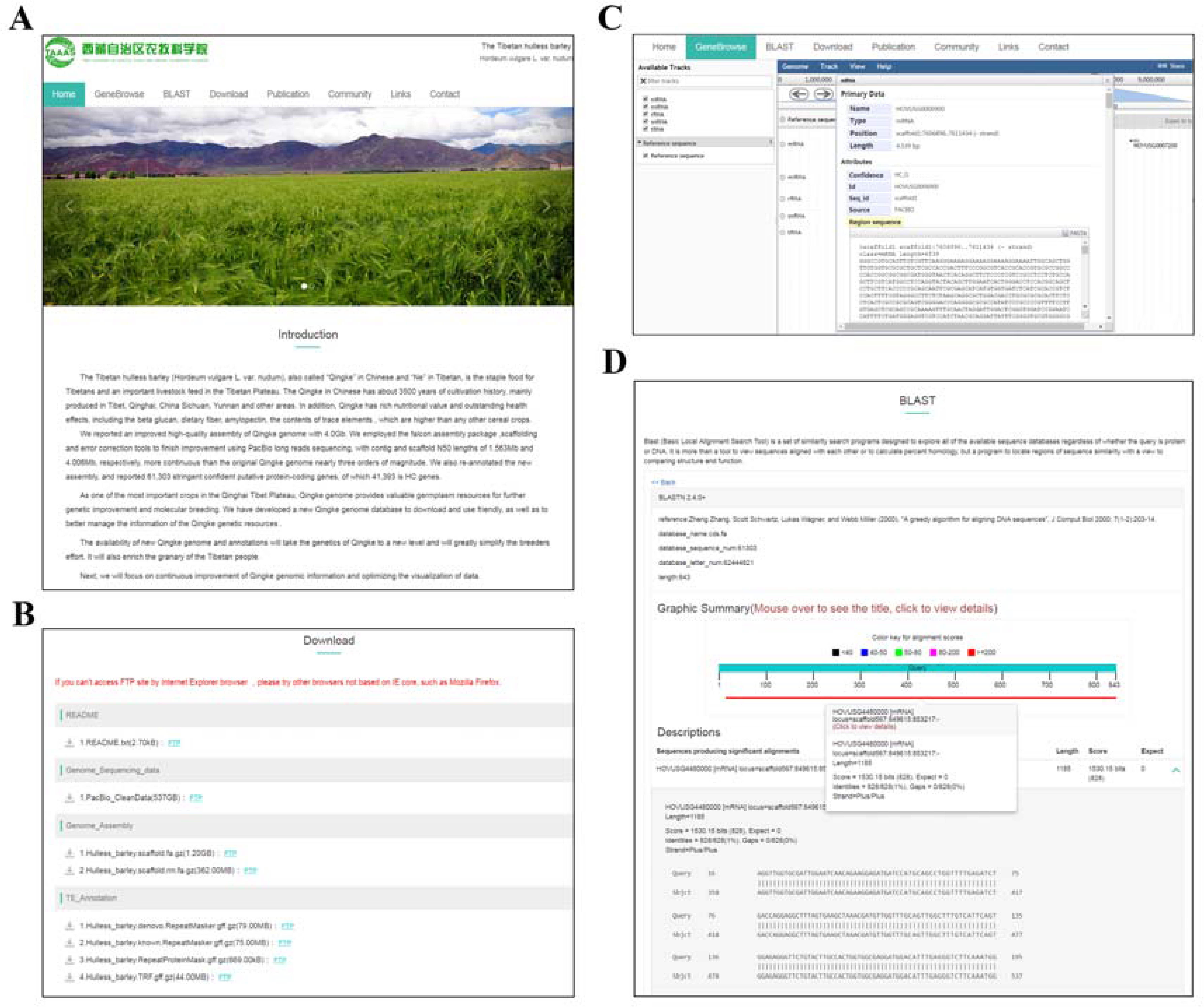
Tibetan hulless barley Genome Database(THBGD). (A),the home page of THBGD;(B), Data download page;(C),gene browse page and (D),sequences BLAST page.

You can access to the THBGD by visiting http://show.genebang.com/project/index

## Discussion

Tibetan hulless barley is a local species of barley and is of great significance to the life and culture of the Tibetan. Its genome and annotation updates help future research on functional genomics, and also help breeders to simplify their work and enrich the genomics resources of barley. The generation of Tibetan hulless barley high-quality reference genomes also demonstrates the superiority of PacBio long reads sequencing technology for larger genome assembly[50]. Although the genome has a long contig, the scaffold is still a bit short at the state-of-art of the current genome assembly, mainly because of the complexity and data volume of the scaffolding mate-pair library is relatively few. In the future, we will use BioNano optical mapping, Hi-C, and akin weapon to complete the genomic construction at the chromosome level[17]. Having an excellent Tibetan hulless barley genomics resources, it will enable geneticists to focus more on functional genomics itself and breeding research.

## Availability of supporting data

Raw genomic sequence reads are available in the NCBI Sequence Read Archive under project number **SRS3725794**. Genome assembly and annotation file are available in Tibetan hulless barley Genome Database (THBGD) or contact the correspondence author. Supplementary files can be downloaded from the GigaScience Website.

## Abbreviations

BLAST: Basic Local Alignment Search Tool
BUSCO: Benchmarking Universal Single-Copy Orthologs
QV: quality value
PacBio: Pacifc Biosciences
RNA-seq: RNA sequencing
NGS: Next generation sequencing
TGS: Third generation sequencing
THBGD: Tibetan hulless barley Genome Database

## Conflicts of Interest

The authors declare no conflict of interest.

## Acknowledgments

This research was supported by the following funding sources: the Tibet Autonomous Region Financial Special Fund 2017CZZX001, 2017CZZX002, XZNKY- 2018-C-021.

We acknowledge (1) Zuoyi Jian, genome assembly executor; (2) Hao Tang, genome annotation pipeline writer; (3) Zhen Zeng, repetitive annotation executor and (4) NextOmics and BGI-Shenzhen for sequencing supports.

Shan Li, Zhenhua Zhuang and Jie Li also contributed to this Paper.

## Author Contributions

Project coordination: X.Z.,X.L.,Y.W.,L.B. and N.T.(Leader).

Sampling and experimentation: X.Z.,Y.W.,H.Y.,Q.X.,S.Z.,Z.W.,W.Q.,Y.B.,J.D.,M.Y..

Genome assembly and Genome annotation guidance and performance:

X.L.(Co-leader) and his team.

Data analysis: T.X., X.Z.,Y.W.,H.Y.,Q.X.,S.Z.,Z.W.,W.Q.,Y.B.,J.D.,M.Y..

Genome database construction: T.X. and his team

Data submission: T.X.

Writing: X.L.(Co-leader).

All authors read and commented on the manuscript.

---

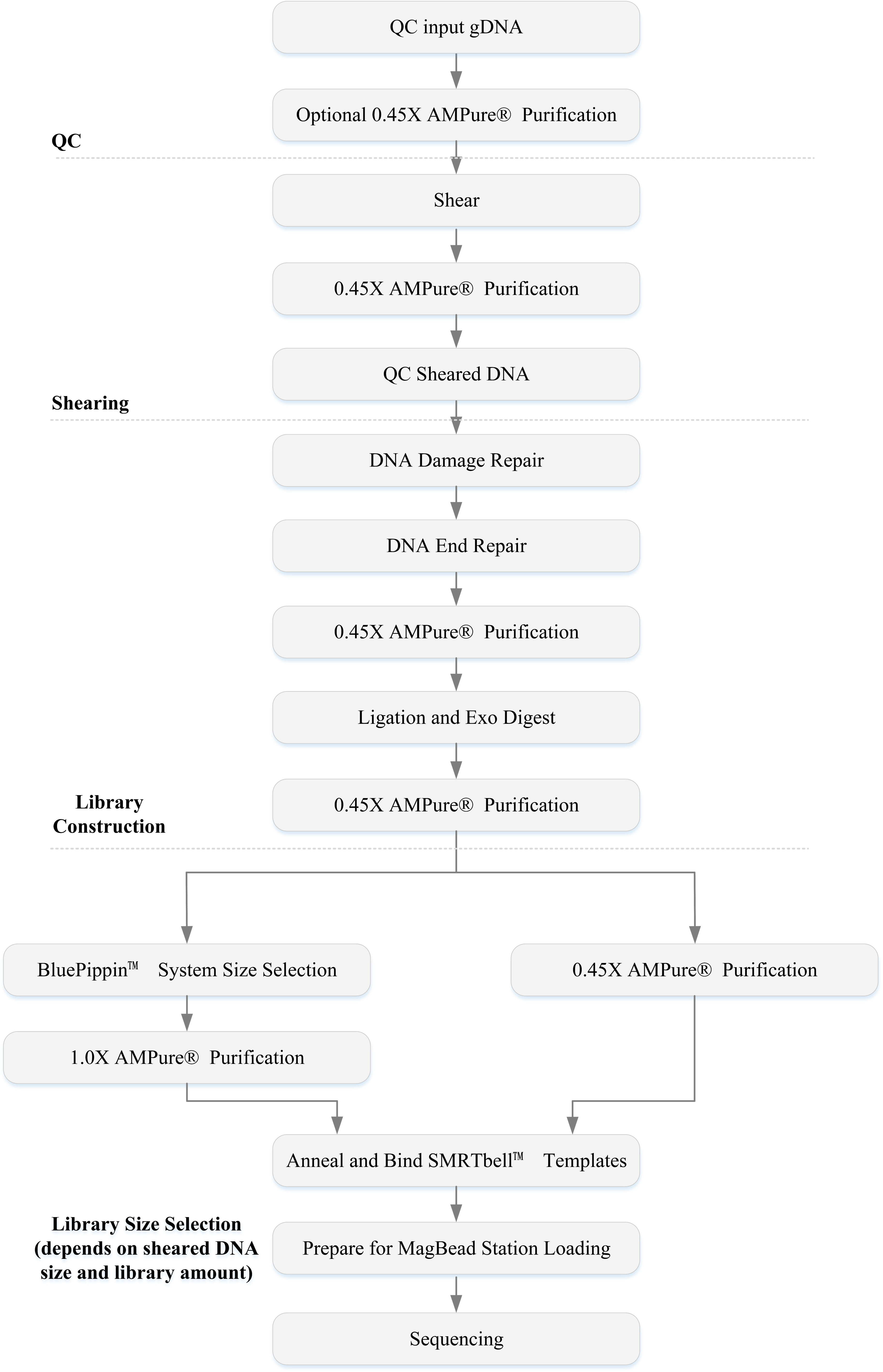

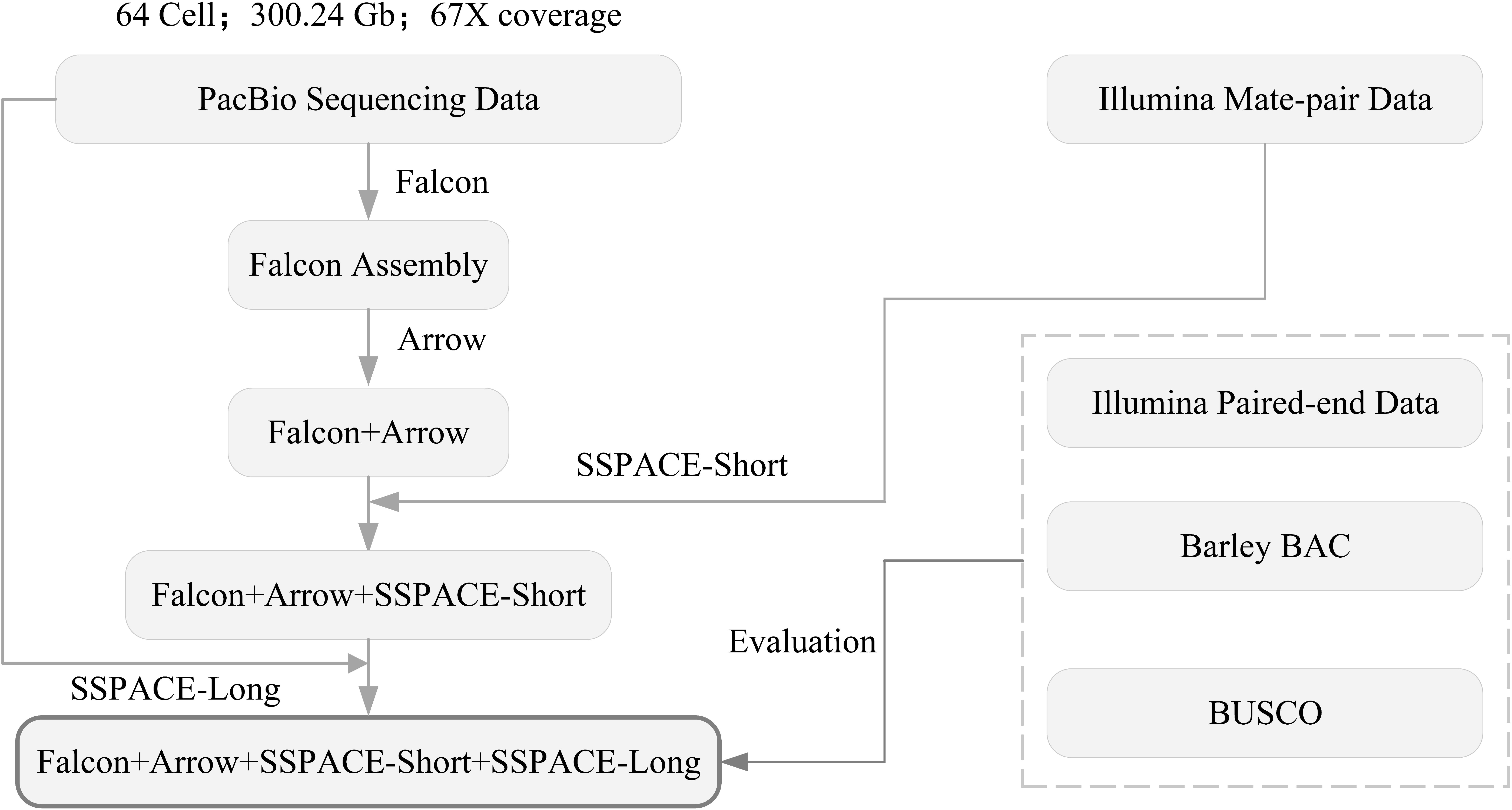

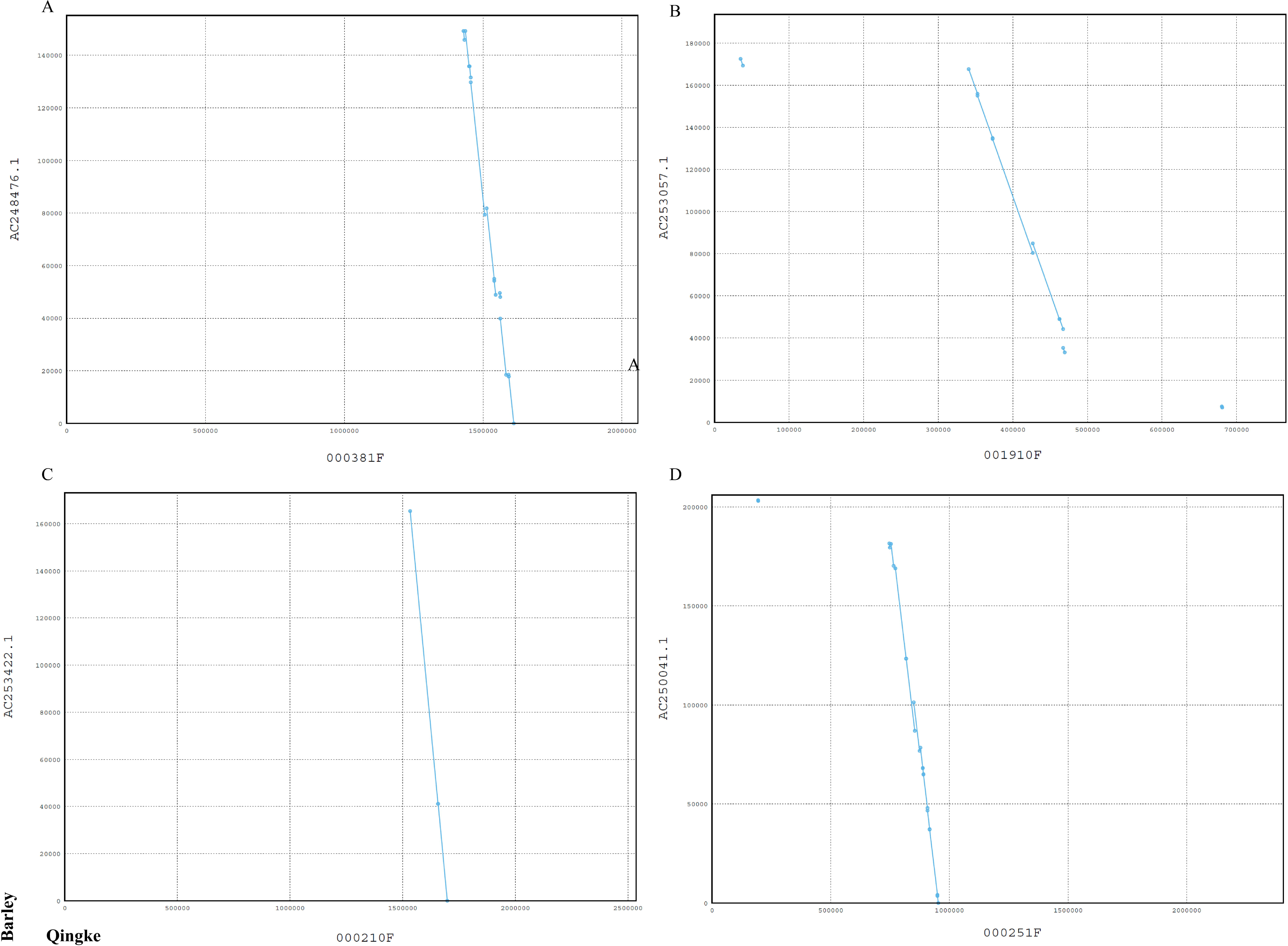

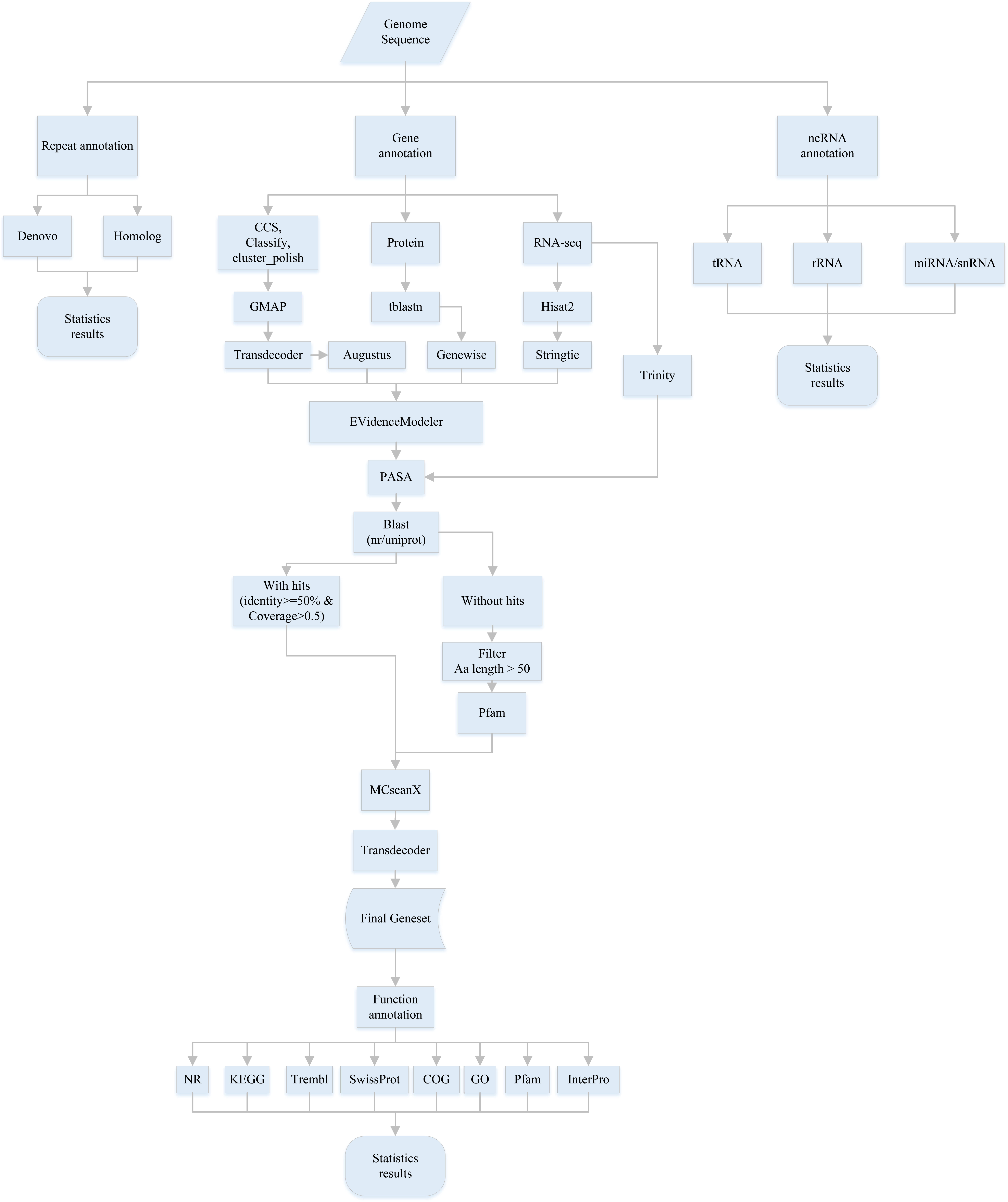

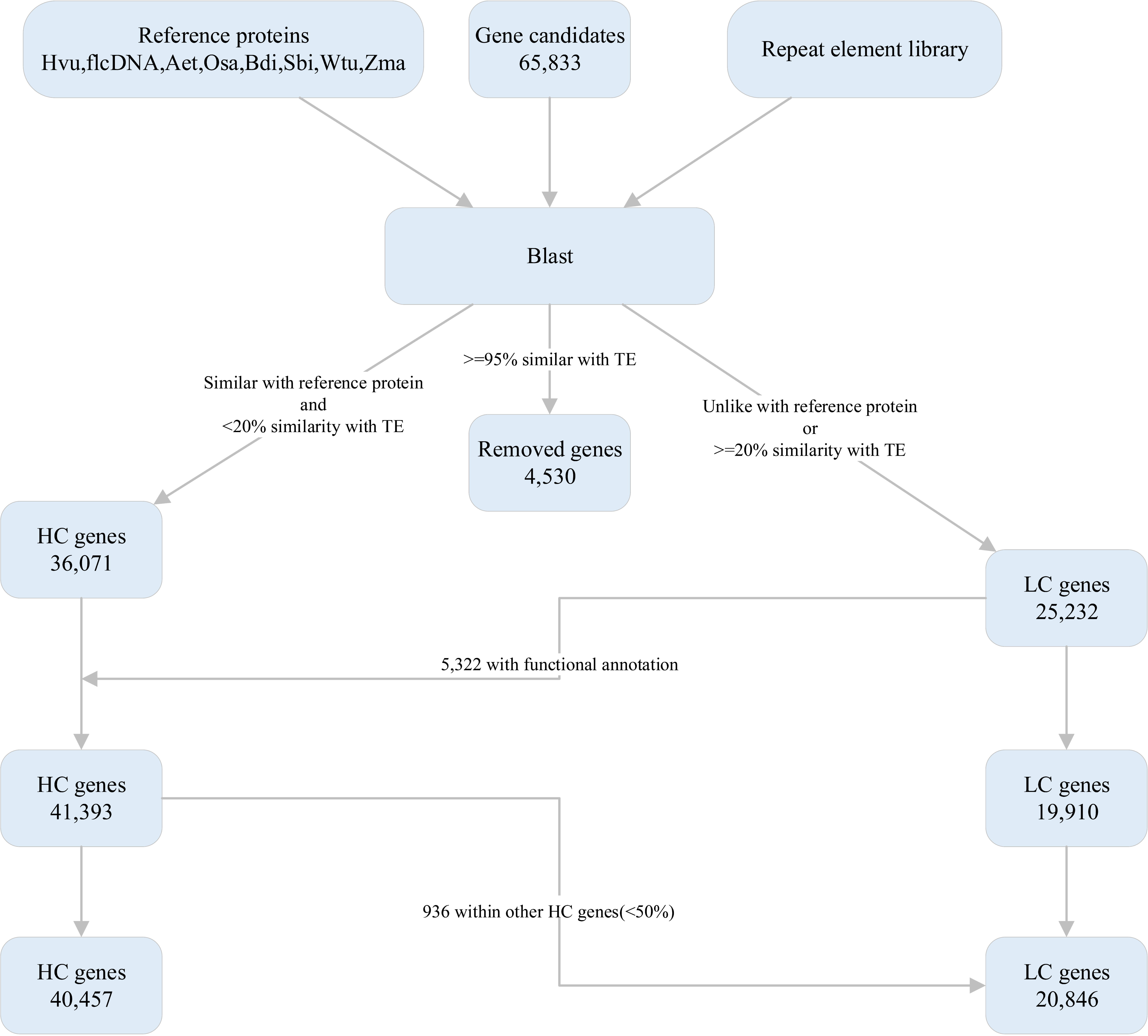

